# Dynamical asymmetry exposes 2019-nCoV prefusion spike

**DOI:** 10.1101/2020.04.20.052290

**Authors:** Susmita Roy

## Abstract

The novel coronavirus (2019-nCoV) spike protein is a smart molecular machine that instigates the entry of coronavirus to the host cell causing the COVID-19 pandemic. In this study, a structural-topology based model Hamiltonian of C3 symmetric trimeric spike is developed to explore its complete conformational energy landscape using molecular dynamic simulations. The study finds 2019-nCoV to adopt a unique strategy by undertaking a dynamic conformational asymmetry induced by a few unique inter-chain interactions. This results in two prevalent asymmetric structures of spike where one or two spike heads lifted up undergoing a dynamic transition likely to enhance rapid recognition of the host-cell receptor turning on its high-infectivity. The crucial interactions identified in this study are anticipated to potentially affect the efficacy of therapeutic targets.

**One Sentence Summary:** Inter-chain-interaction driven rapid symmetry breaking strategy adopted by the prefusion trimeric spike protein likely to make 2019-nCoV highly infective.

---

We are in the midst of a global catastrophic situation due to coronavirus outbreak where every day the global death toll is biting the past count. While history has witnessed past pandemics (*1–3*), 2019 Novel Coronavirus (2019-nCoV) trends to outcompete them all by its rapid transmission in a short period to win the crown. As the name ‘*corona*’ (in Latin, it means crown), the accused of the outbreak makes use of a ‘crown-shaped’ molecular machine, the trimeric spike-protein to drive the virus entry into the host cells. 2019-nCoV is the newest addition of betacoronavirus genus (*4*). While the sequence and structural similarity to the severe acute respiratory syndrome coronavirus spike (SARS-CoV S) repute the identity of 2019-nCoV spike as SARS-CoV-2 S (*5, 6*), a recent study reported that the SARS-CoV-2 S has 10-20 fold higher affinity to human angiotensin-converting enzyme 2 (ACE2) receptor than that of SARS-CoV S (*7*).

The large ectodomain of the S glycoprotein of the coronavirus uses the S1 subunit for receptor binding and the trimeric S2 stalk for host-cell membrane fusion (Fig. 1A) (*8, 9*). In SAR-CoV-2 S glycoprotein, the β-sheet-rich S1 subunit comprises an N terminal domain (NTD) and a receptor-binding domain (RBD) towards its C-terminus (CTD) (Fig. 1B). So far, static structural characterization reported that the RBD of S1 has an intrinsic hinge-like conformational movement that generates the ‘up’ and ‘down’ conformations (*7, 8, 10*). Other betacoronaviruses, like SARS-CoV, MERS-CoV and distantly related alphacoronavirus porcine epidemic diarrhea virus (PEDV) also have this apparently stochastic RBD movement (*11, 12*). The combination of RBD up-down rearrangement may lead each S1-head of the trimeric prefusion spike protein of coronavirus to adopt different possible conformations: (i) 3down, (ii) 1up-2down, (iii) 2up-1down, and (iv) 3up (Fig. 1C). Among them 3down, 3up are symmetric conformers and 1up-2down, 2up-1down are asymmetric conformers. Single-particle cryo-electron microscopy (Cryo-EM) determined few such symmetric and asymmetric structures referred to as the receptor-binding inactive state and receptor-binding active state, respectively (*8*). The asymmetric structure where one of the RBDs rotates up was thought to be less stable for SARS-CoV S (*10*). In comparison, the recent Cryo-EM study found three RBDs in 1up-2down conformation as a predominant arrangement in the prefusion state of 2019-nCoV S trimer (*7*). This arrangement apparently appears legitimate for SARS-CoV-2 S in order to explain the higher affinity of 1up-2down for ACE2 receptor than that of SARS-CoV S. However, we cannot rule out the possibility of 2up-1down conformation as a functional state, which may provide even stronger binding with ACE2 considering the fact that ACE2 is a dimeric receptor (*9, 13*). This hypothesis is consistent with a recent crystallographic study demonstrating that CR3022, a neutralizing antibody isolated from convalescent SARS patients targets the RBD when at least two RBD on the trimeric spike protein are in the up conformation (*14*). Assembling all these experimental results it is high time to understand the molecular mechanism of S1-head coordination of trimeric SAR-CoV-2 S and to identify important interaction in regulating spike up-down conformations.

**Fig. 1.**
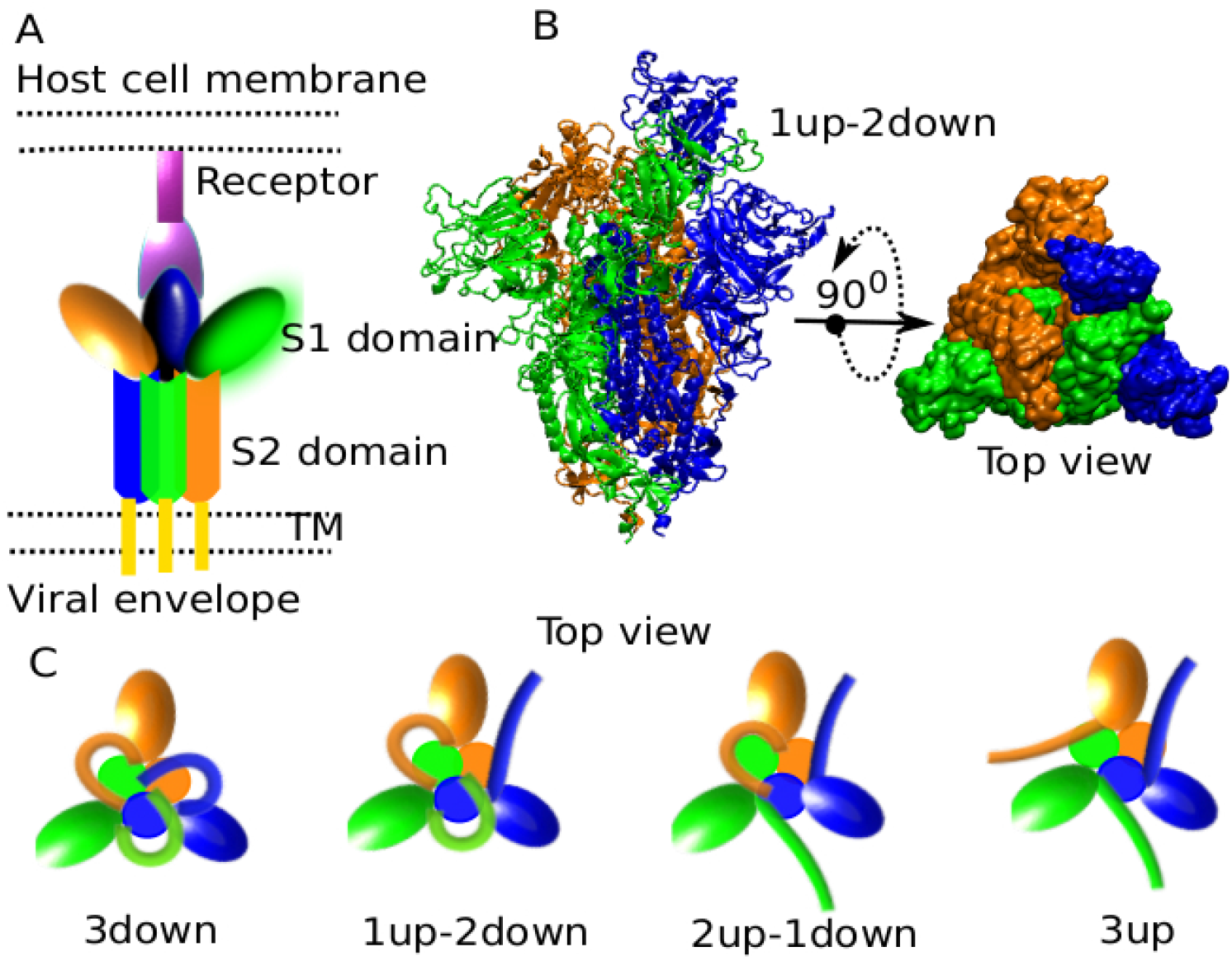
Conformational illustration of coronavirus spike proteins. (A) A schematic of receptor-bound spike protein including the receptor-binding subunit S1, the membrane-fusion subunit S2 of a coronavirus is shown. B. Side and top views of the homo-trimeric structure of SARS-CoV-2 spike protein with one RBD of the S1 subunit head rotated in the up conformation. C. RBD up-down movement expected to lead S1 heads of the trimeric spike protein to attain the following possible conformers: (i) 3 down (ii) 1up-2down (iii) 2up-1down, and (iv) 3up. These are an analogue demonstration of the spike protein top-view where NTDs are represented by colored ovals, RBDs are represented by flexible sticks and S2 domains are represented by filled circles.

A major challenge was simulating the gigantic structure of the full-length trimeric spike, as it is associated with the largescale conformational transition. It is indeed a daunting task to explore the full conformational landscape at an atomic length-scale. To overcome this, a structure-based coarse-grained molecular dynamic simulation approach has been adopted (*15*). The simulation started with a full-length homo-trimeric spike protein structure generated from homology modeling which involves the alignment of a target sequence and a template structure (pdb: 6vsb) (*7, 16*). This also helped to build the missing loops. The domain-specific residue-range for the full-length, trimeric SARS-CoV-2 S is given in Fig. 2A. The S1 head coordination of the trimeric spike is programmed by developing a super-symmetric topology-based modeling framework (Fig. 2B) (described in the Method pipeline in the supplementary material). With this, the molecular machine is ready to swing each of its S1 head between its ‘up’ and ‘down’ conformations (Movie S1, S2). A number of Cryo-EM structures captured the ‘up’ and ‘down’ conformations of the RBD domain of spike proteins of other coronaviruses including SARS-CoV-2 where the S1 subunit undergoes a hinge-like conformational movement prerequisite for receptor binding (Fig. 2C) (*7, 8, 10, 17*). Apart from the hinge-responsive RBD-cleft interaction, in this study, a few inter-chain interactions are found to assist the ‘RBD-up’ and the ‘RBD-down’ conformations (shown in Fig. 2D and 2E, Movie S3). These few interactions are identified to impact the breathing of RBD of SARS-CoV-2 S. This makes the early referred ‘RBD-up/down’ conformations slightly different from the ‘S1-head-up/down’ conformation for trimeric SARS-CoV-2 S as the former is regulated only by intra-chain interactions while the latter is regulated by both intra and inter-chain interactions (Fig. S1). After identifying all these unique intra and inter-chain contacts (*18, 19*) extracted from the corresponding ‘S1-head-up’ and ‘S1-head-down’ conformations, a super-symmetric contact map is generated. This follows the development of a structure-based model Hamiltonian (Materials and Methods in Supplementary) which is based on the energy landscape theory of protein folding (*20–24*). This approach not only potentiates the trimeric spike to adopt C3 symmetric ‘3up’ and ‘3down’ states but also to break the symmetry in a thermodynamically governed way (Fig. S1-S4) (*25, 26*).

**Fig. 2.**
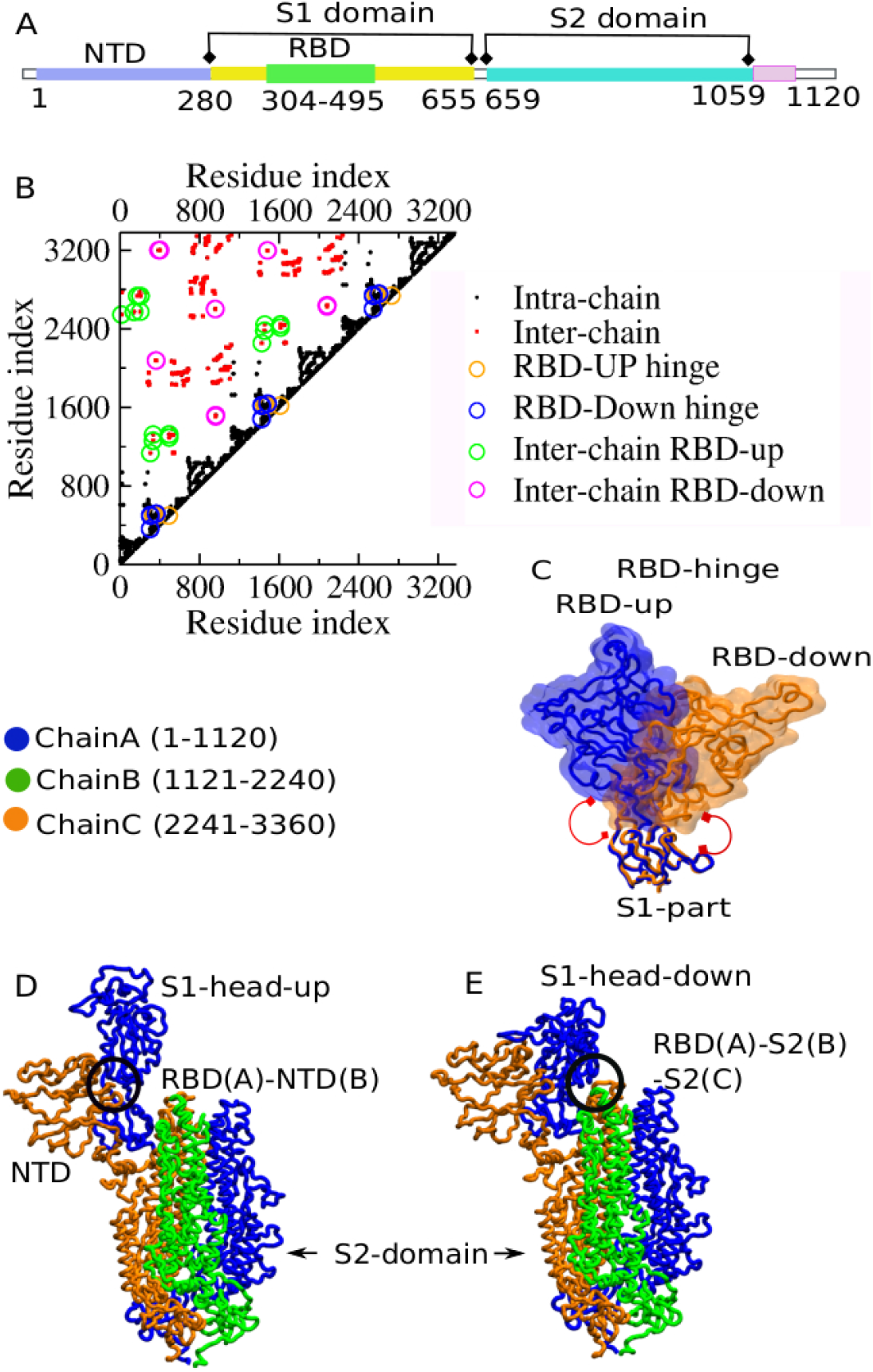
Building a super-symmetric contact map of the homotrimeric SARS-CoV-2 spike protein. A. Amino acid sequence ranges of NTD, RBD, and S2-subunit are only highlighted. B. Residue-residue native contact map identifying unique intra and inter-chain contact-pairs formed by any single monomer in its S1-head up and S1-head down states. C. Within intra-chain contacts, the unique contacts that drive hinge motion leading to RBD-up and RBD-down states are highlighted in the structure, as well as in the contact map. D. Inter-chain unique contacts between RBD and NTD domains upholding the S1-head-up state. E. Inter-chain unique contacts are responsible for connecting the RBD of ChainA with the S2-stalk of ChainB and the S2 stalk of ChainC.

To monitor the transition between the ‘S1-head-up’ and the ‘S1-head-down’ states for each monomer with the trimeric interactions, a large pool of unbiased longtime trajectories generated where multiple occurrences of up and down states for each monomer have been sampled. We employ a reaction coordinate, Q, the fraction of the native contact (*19, 27*) corresponding to the inter-chain contacts associated with the ‘S1-head-up’ and the ‘S1-head-down’ states. A typical trajectory plot of Q extracted from the equilibrium simulation of the trimeric prefusion spike clearly shows the hopping between different conformational states as hypothesized earlier (Fig. 3A). Furthermore, the dynamic transitions between the two major asymmetric states (1up-2down: Q_S1-head-down_≈0.45 and 2up-down: Q_S1-head-down_≈0.15) are evident in the Q-trajectory. Analysis of all the simulations yields the 2-D free energy landscape of the trimeric spike protein of SARS-CoV-2 (Fig 3B) with its all possible conformations. The conformations corresponding to the minima of the free energy landscape are shown in Fig.3C. The temperature dependence of conformational transition indicates that the configurational entropy and enthalpy compensation results in the enhanced population of the asymmetric 1up-2down to 2up-1down conformations (Fig S4). While the predominant population of the 1up-2down state is consistent with the recent Cryo-EM data, (*7*) (Movie S1, S2) the other asymmetric structure (2up-1down) emerges as a best binding epitope for CR3022 (an antibody collected from convalescent SARS patients) according to a recent antibody recognition study of SARS-CoV-2 S (*14*).

**Fig. 3.**
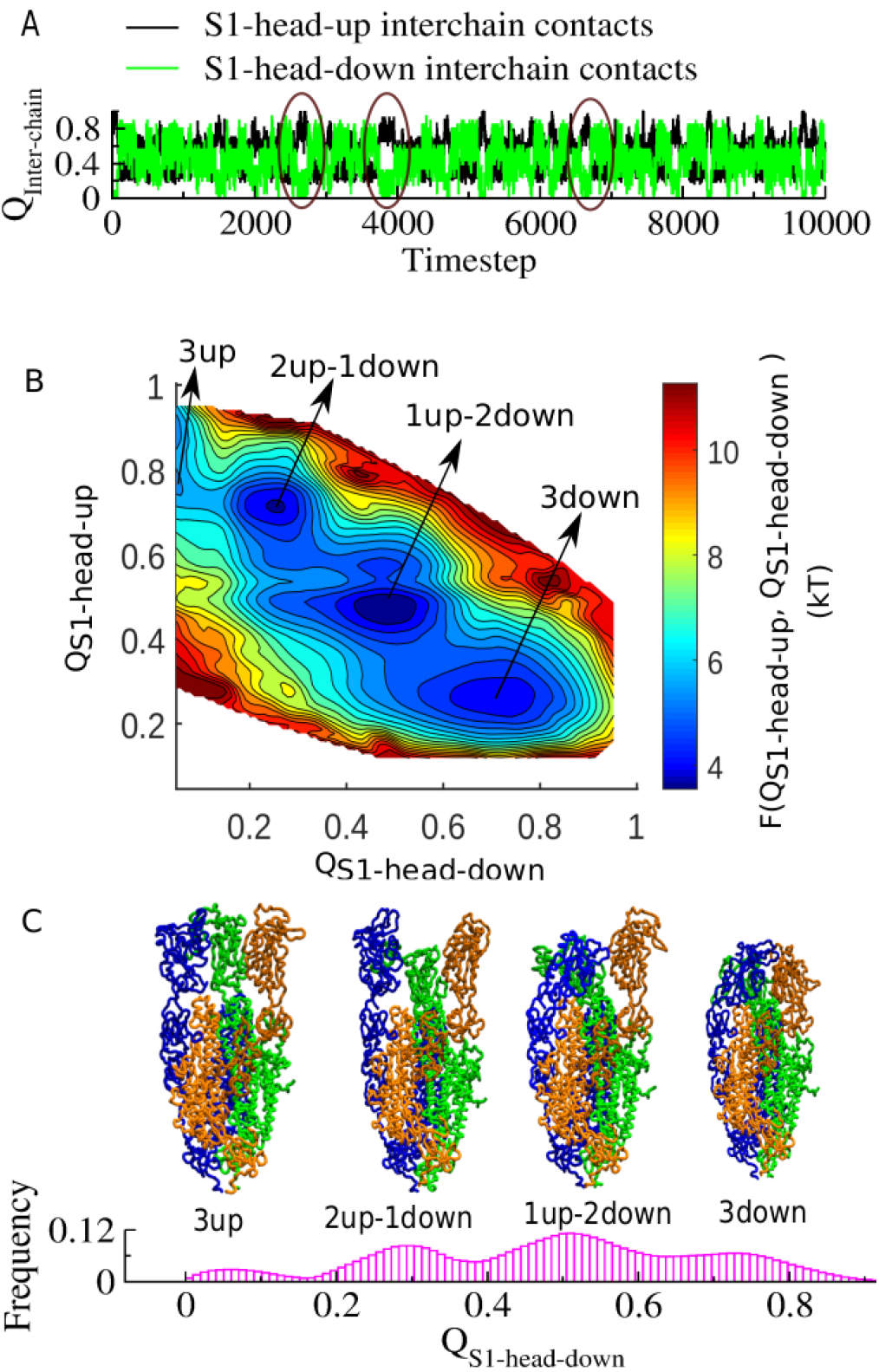
Conformational transition of SARS-CoV-2 spike protein in its prefused state. A. The fraction of native contact (Q) dynamics counting inter-chains contact-pairs formed in the S1-head-up state and the S1-head-down state. B. A two-dimensional free energy landscape of conformational transition as a function of inter-chain contacts supporting S1-head-down (x-axis) and S1-head-up state (y-axis) explores all possible conformations. C. The representative structure corresponding to each minimum of the free energy landscape is designated as follows: (i) 3up, (ii) 2up-1down, (iii) 1up-2down, and (iv) 3down state (as shown in the one-dimension population distribution plot).

In this study, sequence and interaction level (Fig. S5, Fig. S1) comparison has been made over the Cryo-EM structure of SARS-CoV-2 S (pdb: 6vsb), SARS-CoV S (pdb:5×5b) and MERS-CoV S (pdb: 5×5f) (*7, 12*). This comparison results that SARS-CoV-2 S has NTD-RBD domain association where a proline residue of ChainA forms CH-п type interaction with the tyrosine residue (*28*) and hydrophobic interaction with another proline of ChainB (Fig. 4A, Fig. S1). Inter-chain proline-proline distance measurement shows that the corresponding RBD-NTD domains are far away in the case of SARS-CoV S (Fig. 4B) and further away in the case of MERS-CoVS (Fig. 4C). This measurement involves their respective Cryo-EM structures. Despite the relatively high degree of sequence similarity between the SARS-CoV-2 S and the SARS-CoV S and also with the spike protein from the bat coronavirus RaTG13, a single histidine residue at the relevant RBD-NTD domain interface is found unique in the vase of SARS-CoV-2 S (Fig. S5) (*29*). The imidazole ring of histidine is pointing towards the hydrophobic assembly of aforesaid proline-tyrosine in the juxtaposition of the RBD-S1 hinge region. Such inter-chain RBD-NTD connection is thus found to impact the RBD hinge interaction by upregulating more RBD-up conformation (Fig. 4D). In the absence of such inter-chain interaction, the RBD mostly stays in the down conformation allowing RBD to break the symmetry rarely in a stochastic manner (Fig. 4E). The absence of inter-chain RBD-NTD connection also appears to impact the SARS-CoV RBD hinge interaction. Here, the opening of RBD-S1 cleft is significantly less than that of SARS-CoV-2 S in their respective S1-head-up state (Fig. S6). The assistance from the inter-chain RBD-S2-stalk related interfacial contacts are also found to modulate the population dynamics of RBD-down conformation (Fig. S7). The influence of this inter-chain RBD-S2-stalk interaction has also been observed in an early Cryo-EM analyses where two proline mutations at the top of S2 stalk (inferring RBD-S2 inter-chain connection) helped to stabilize the ‘up’ conformers of SARS-CoV S (*30*).

**Fig. 4:**
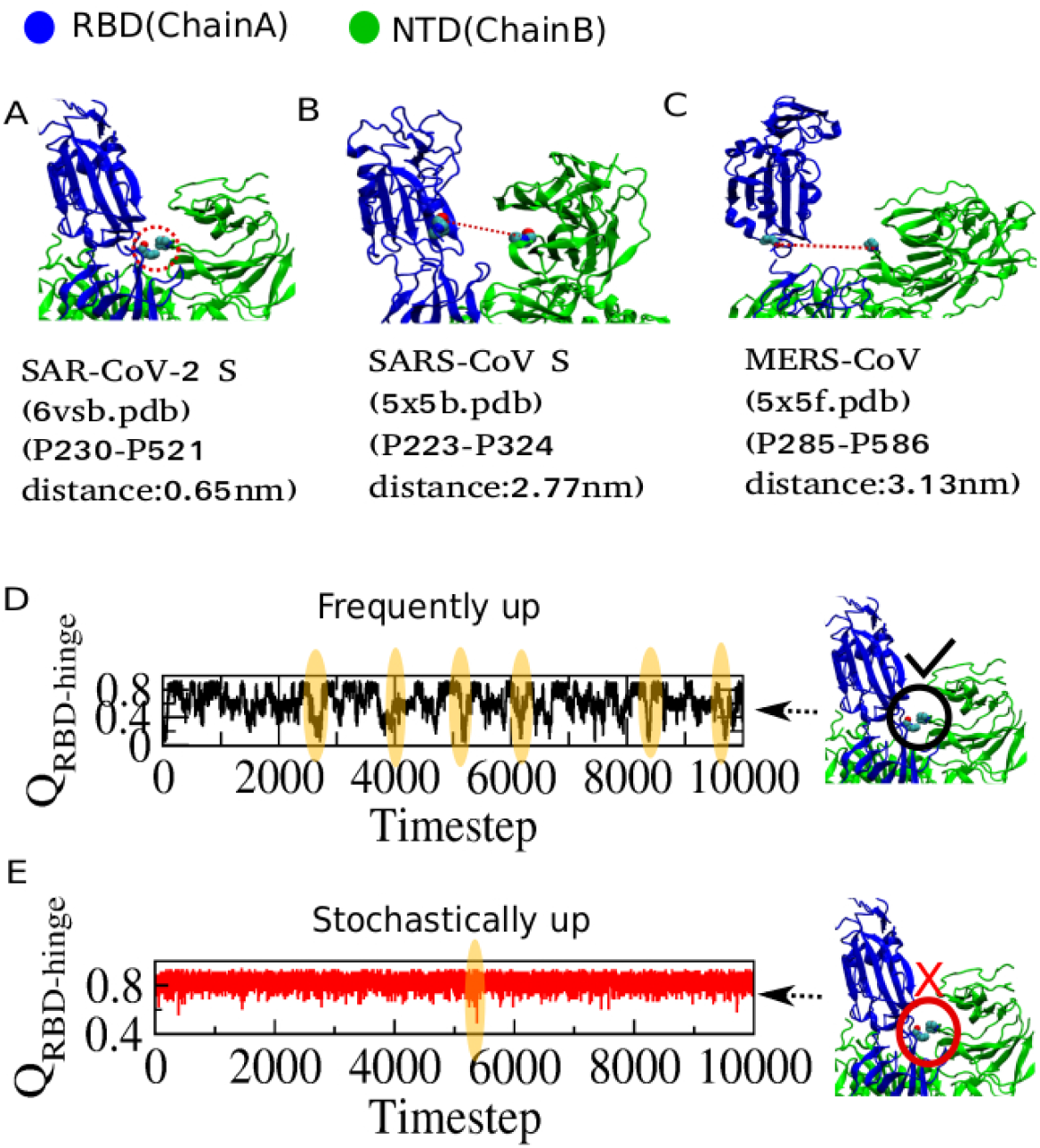
Interactions that make SARS-CoV-2 spike different from SARS-CoV and MERS-CoV spike. A. Unique inter-chain interactions formed by RBD of one chain with NTD of the adjacent chain stabilizing the S1-head-up conformation in SARS-CoV-2 S (pdb:6vsb). Inter-chain domain closure is analyzed by inter-chain proline-proline distance measurement. The same distance measured for the following spikes: B. SARS-CoV spike (pdb:5×5b) and C. MERS-CoV spike (pdb: 5×5f). D. RBD up-down hinge dynamics triggered by inter-chain RBD-NTD domain interaction. E. In the absence of RBD-NTD inter-chain interaction, the hinge motion of RBD is hindered by populating more ‘RBD-down’ conformations and allows to sample ‘RBD-up’ conformation only rarely in a stochastic manner.

The synergy between internal RBD-hinge interactions and inter-chain interactions allows trimeric SARS-CoV-2 S to adopt a unique dynamical feature than other corona-virus spikes. It appears that the inter-chain interactions driven rapid symmetry breaking strategy potentiates this spike machine to turn on its high-infectivity. The energy landscape framework used in this study indeed helps to unify and compare different spike protein interactions present in other coronaviruses. While in the current situation to develop diagnostics and antiviral therapies are of utmost priority, the present structure-based model derived information at the microscopic interaction level might provide deep insight to design effective decoys or antibodies to fight against 2019-nCoV infection.

## Acknowledgments

Author thanks DIRAC supercomputing facility at IISER-Kolkata for computational support.

## Author contributions

S.R. designed the research, built the structure-based model of spike protein, performed simulations, analyzed results and wrote the manuscript.

## Data and materials availability

All data and codes used in the analysis are available from the corresponding author to any researcher for purposes of reproducing or extending the analysis under a material transfer agreement with IISER-Kolkata, India.

## Supplementary Materials

### Materials and Methods

#### Method pipeline of building a super-symmetric contact map of SARS-CoV-2 prefusion spike protein

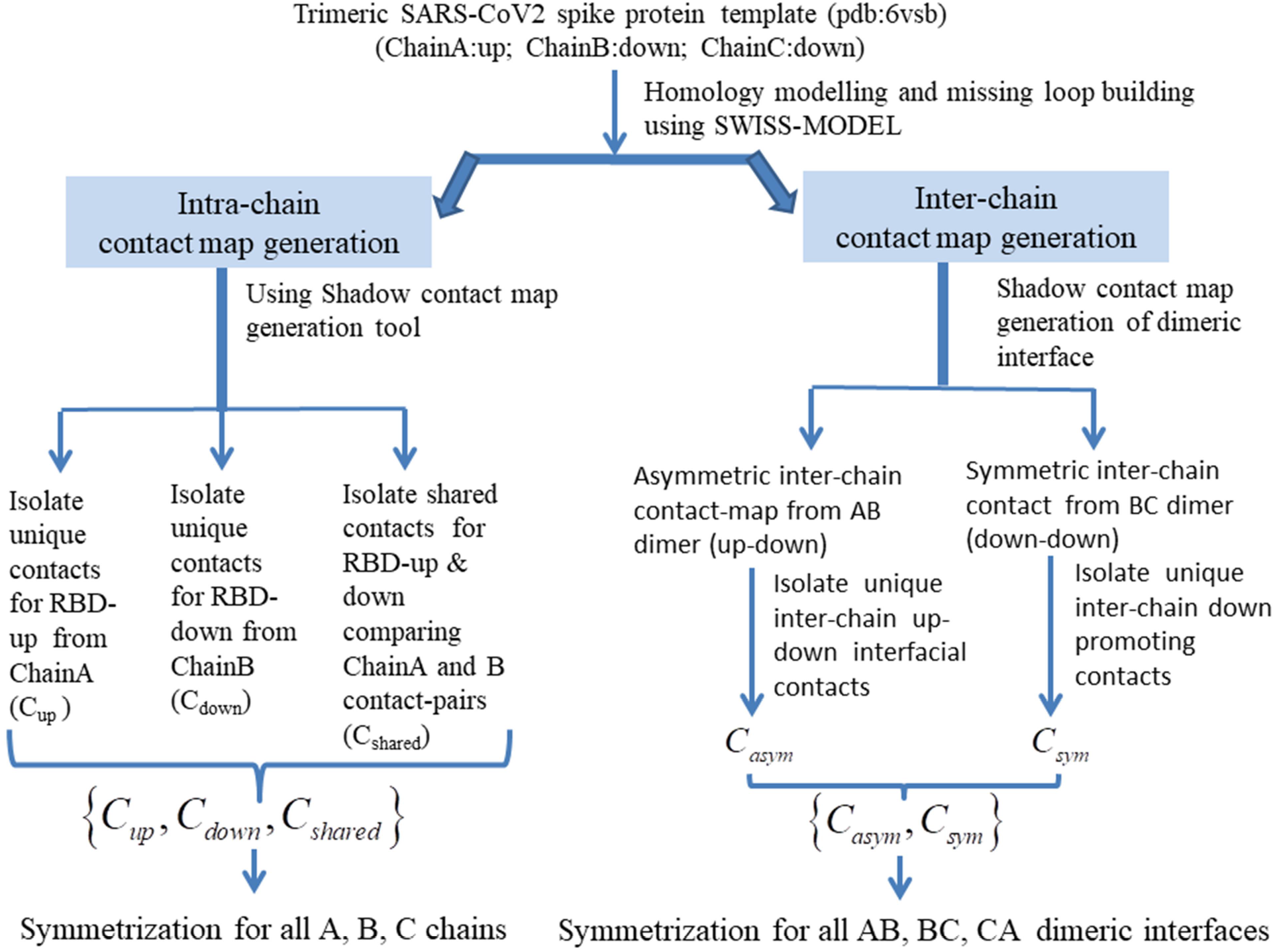

##### Super-symmetric contact map generation method

Coarse-grained structure-based simulations have been performed for full-length trimetric SARS-CoV-2 spike protein. The structure-based Hamiltonians for different simulations were derived after processing the recent Cryo-EM structure (pdb:6vsb) thorough the Swiss Model to complete missing loops present in the structure (*7, 16*). This generates a homo-trimeric SARS-CoV-2 spike where this initial structure has important components in terms of intra and inter-chain contacts (interaction) leading to an ‘S1-head-up’ and an ‘S1-head-down’ conformation for each protomer. In this prevalent trimeric variant, only one monomer adopts ‘S1-head-up’ and the same of the other two adopts the ‘S1-head-down’ conformation. Few characteristic intra-chain contacts cause the receptor-binding domain to perform a hinge-motion resulting ‘RBD-up’ and ‘RBD-down’ conformations for each monomer. Intra-chain contact-pair list is generated separately from ChainA which is a representative of the RBD-up conformation as well as from ChainB which is a representative of the RBD-down conformation. To avoid over-counting the contact-pairs of ChainA and ChainB are compared. Only unique contacts related to RBD-up (*C*_up_) and RBD-down (*C*_*down*_) conformations and a set of shared contacts (*C*_shared_) are considered as shown in the pipeline method. This set of {*C*_up_, *C*_down_, *C*_shared_} is symmetrically distributed over all chains making each of them dynamically capable of sampling both RBD-up and RBD-down conformations driven by *C*_intra_ as defined in the pipeline method. Contact calculation is performed using the Shadow criterion (*19*).

Interesting components are inter-chain contacts residing at the interface of the dimer. Now, two categories of interactive dimeric interfaces are there: asymmetric-dimer interface and symmetric-dimer interface. ChainA (S1-head-up) and the adjacent ChainB (S1-head-down) represent an asymmetric dimer unit. Similarly, ChainB (S1-head-down) and the adjacent ChainC (S1-head-down) represent a symmetric dimer unit. At the asymmetric-dimer interface, the RBD-domain of ChainA forms a few unique contacts with the NTD domain of the adjacent ChainB as shown in Fig. 2D mostly ensuring the S1-head-up arrangement. This asymmetric-dimer also includes a relatively remote RBD(B)-S2(A) contacts (Fig. 2E) that can bend down the S1-head of a chain only partially. All these dimeric interfacial contacts are unique and identified as *C*_*AB*_. Similarly, in the asymmetric-dimer interface, the RBD-domain of ChainB forms a few unique contacts with the S2 domain of the adjacent ChainC as shown in Fig. 2E. These contacts can bend down the S1-head of a chain only partially. In trimeric spike, RBD(B)-S2(A) is equivalent to RBD(A)-S2(C) and RBD(C)-S2(B) following the cyclic rule. Our analysis shows that the complete bent-down of S1-head resulting in the S1-head-down conformation is collectively determined by RBD(A)-S2(B), RBD(A)-S2(C) type of contact elements along with intra-chain hinge contact. Finally, interface-related contact set{*C*_*AB*_, *C*_*BC*_} has been cycled over all the interfaces making each of interfaces dynamically capable of inducing S1-head movement.

##### Developing a structure-based Hamiltonian of trimeric spike protein simulation

A structure-based Hamiltonian of trimeric spike protein for SAR-CoV2 is derived using the super-symmetric contact map. In the current structure-based model amino acids are represented by single beads at the location of the C-α atom (*15, 25, 31, 32*). The coarse-grained structure-based model, a well-established model, comprehends a novel way to investigate the mechanisms associated with protein folding and function (*20–22, 24, 33–38*). In the current context of decoding virus entry mechanism, this model successfully characterized Class-I viral fusion protein dynamics including conformational rearrangement of a viral surface glycoprotein, influenza hemagglutinin (HA) during its prefusion and postfusion states (*26, 39*).

As described in the pipeline method, the complete Hamiltonian comprises of two terms: *H*_intra_, *H*_inter_ The complete Hamiltonian as a function of a set of position coordinates 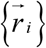 has the following simple form:

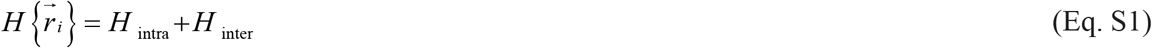

Where *H*_intra_ contains three symmetric terms applied over chain A, Chain B and Chain C to maintain their internal local and non-local interactions.

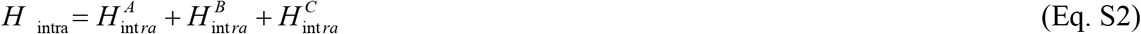

The local part of 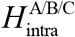 includes harmonic potentials that restrain bonds (*r*), angles (*θ*). Dihedral angles (*ϕ*) are treated with a cosine term, as shown in Eq. S4. The initial geometric parameters 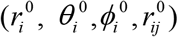 are obtained from the initial trimeric spike structure as shown in Fig 1B.

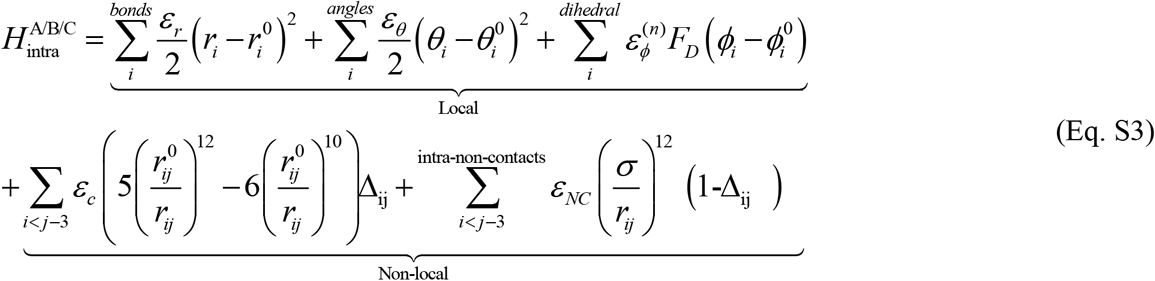

Where,

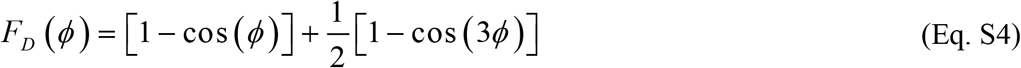

And,

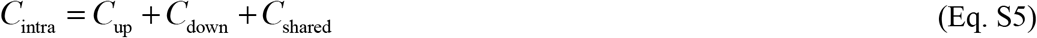

The first non-local term of the Hamiltonian used in 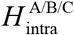 represents non-bonded interaction potential in the form of 10–12 Lennard-Jones potential that is used to describe the interactions that stabilize the native contacts(*15*). A native contact is defined for a pair of residues (*i* and *j*) present in the native state using shadow criteria and when (*i*−*j*)>3. Δ_*ij*_ is defined in such a way that if any *i* and *j* residues belong to *C*_intra_, Δ_*ij*_ = 1 turning on 10–12 Lennard-Jones potential; otherwise Δ_*ij*_ = 0. For all non-native pairs for which Δ_*ij*_ = 0, a repulsive potential with σ = 4Å is used. All the interaction coefficients used in this potential are given in **Table S1**.

As described in the method pipeline, *H*_inter_ will include only the non-local inter-chain contacts residing at the interface of the dimer which comprises of accounting for asymmetric-dimer interfacial contacts *C_asym_* and symmetric-dimer interfacial contacts *C_sym_* those are symmetrized for all AB, BC, CA dimeric interfaces.

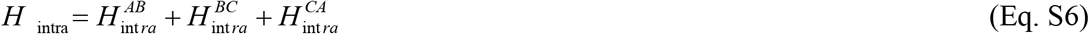

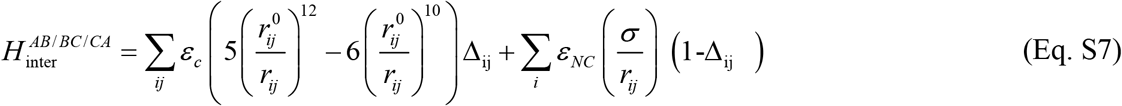

Similar to our early approach, Δ_*ij*_ is such defined that if any *i* and *j* residues belong to *C*_inter_, Δ_*ij*_ = 1, turning on 10–12 Lennard-Jones potential; otherwise Δ_*ij*_ = 0. Here, *C*_inter_ = *C*_*AB*_ + *C*_*BC*_ + *C*_*CA*_

##### Equilibrium Simulation Details

To begin every simulation an initial structure is energetically minimized under the structure-based Hamiltonian using the steepest descent algorithm. Atomic coordinates of the energy minimized structure have been evolved using Langevin dynamics with a time step of 0.0005 *τ*_*R*_. We used an underdamped condition for rapid sampling(*40*). For explicit particles, reduced mass of 1 *μ*_*R*_ and a drag coefficient 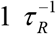 are used to efficiently sample the conformational space(*15, 41*) where 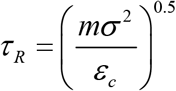 Standard structure-based models implemented in Gromacs (SMOG) were used for all the simulations (*18*).

**Table S1:** Values of parameter set used for current structure-based spike simulation

| **<u>Reduced parameter set</u>** |  |
| --- | --- |
| $\mu_R$ | 1 amu |
| $\epsilon_R$ | 1 kJ/mol |
| $T_R$ | 120 K |
| **<u>SBM potential</u>** |  |
| $\epsilon_r$ | $2 \times 10^2 \epsilon_R / \text{nm}^2$ |
| $\epsilon_\theta$ | $20.0 \epsilon_R / \text{rad}^2$ |
| $\epsilon_\varphi (1)$ | $1.0 \epsilon_R$ |
| $\epsilon_\varphi (3)$ | $0.5 \epsilon_R$ |
| $\sigma$ | 0.4 nm |
| $\epsilon_{NC}$ | 1.0 kJ/mol |
| $\epsilon_C$ | 1.0 kJ/mol |

###### Temperature-Dependent Simulations and Analysis

All temperatures mentioned here are in reduced units. Temperature dependence of the conformational transition has been performed over several temperatures. Three representative reduced temperature-dependent (T*=0.50T_R_, T*=0.58T_R_, and T*=0.83T_R_) analyses are shown for clarity in Fig. S4. Population distribution as a function of the fraction of native inter-chain contacts formed in the S1-head-down state is monitored over these temperatures. Four states emerge as indicated in Fig. 3B and Fig. S4. As the temperature increases the population shifts more towards the S1-head-up state. At T*=0.58T_R_ the population of 1up-2down state appears as a predominant population in the conformational landscape which correlates well with the recent Cryo-EM data (*7*). We have performed all our simulations being consistent with this selected temperature. The RMSD analyses ensure the correctness of the simulation progress and the emergence of the correct structure (Fig. S3).

The population shifts more towards the S1-head-up state conformations as the temperature increases. It suggests that the S1-head-up states are more dynamic and entropically stable. Note that the dynamical transition between 1up-2down and 2up-1down states may tolerate a wide range of temperatures by a population shift mechanism. So far, we have examined that it tolerates the temperature range from T*=0.50T_R_ to T*=1.67T_R_. Temperature dependence of RBD hinge motion has also been studied (Fig. S4). Population distribution as a function of the fraction of native intra-chain hinge-region contacts formed by the RBD at different temperatures has been monitored. A bimodal distribution reflects the population of the ‘RBD-up’ and ‘RBD-down’ states for any individual chain being in trimeric spike. As temperature increases, the RBD-up states start to enhance their populations.

##### Free energy calculation

In a system, if a state “A” described by its reaction coordinate, *X_A_* (which in our case is the fraction of native contact) is separated from another state “B” described by its reaction coordinate, *X_B_*, by a finite barrier, the free energy of transition from A to B can be expressed as,

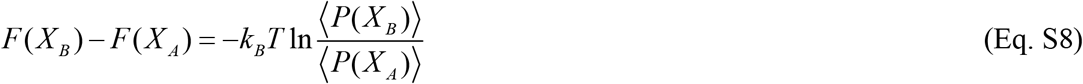

where, 〈*P*(*X_B_*) 〉 is the probability to find the system in state B at the reaction coordinate, Q_B_. the same holds for 〈*P*(*X_A_*) 〉. From a finite set of unbiased simulations of trimeric spike protein, a complete thermodynamic description is obtained. Probability distributions are obtained by sampling the configurational space running 50 Molecular Dynamics simulation sets.

---

**Fig. S1:**
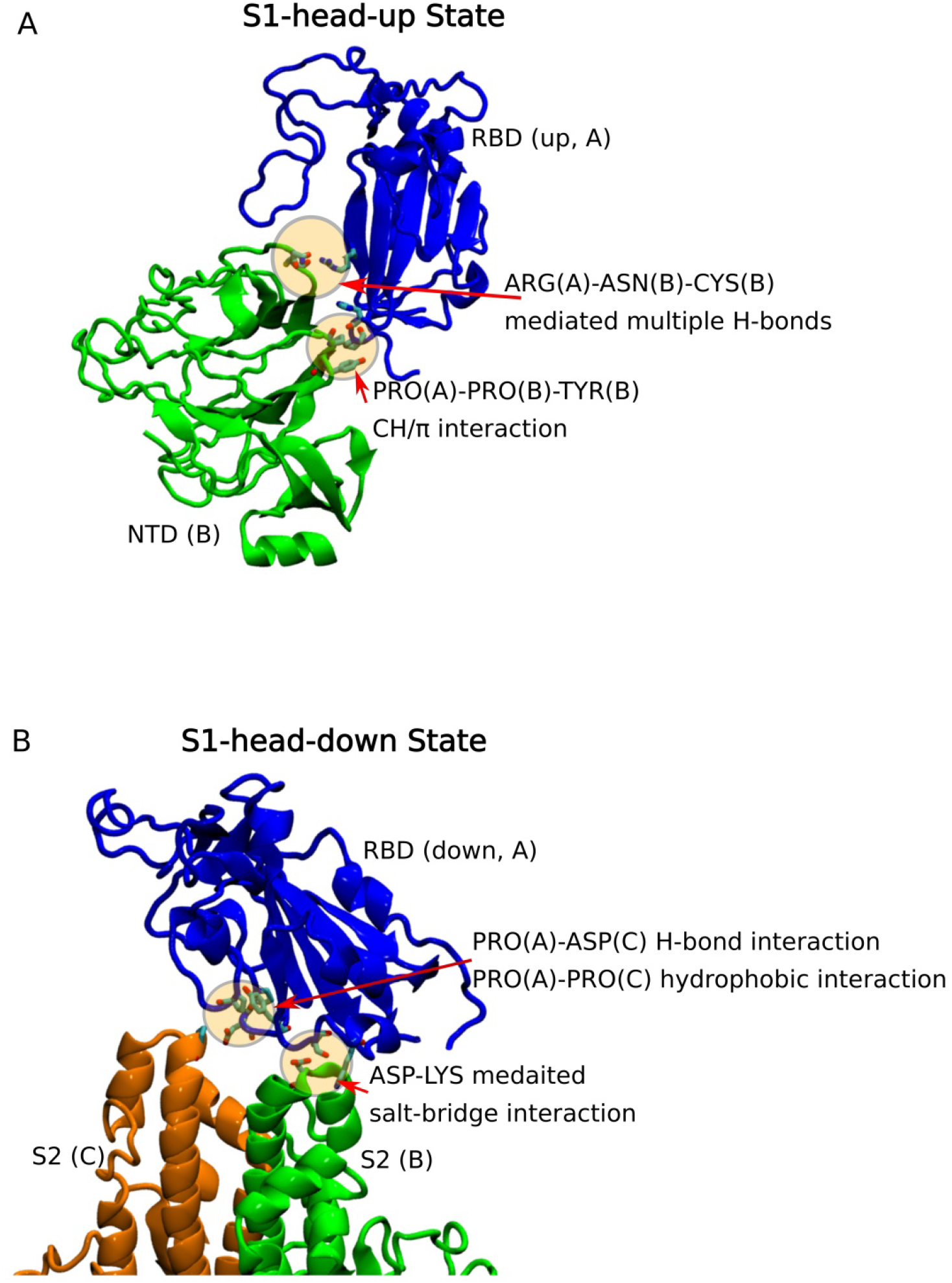
Inter-chain interaction from the ‘S1-head-up’ and the ‘S1-head-down’ states of SARS-CoV-2 spike. A. Inter-chain RBD-NTD domain closure in the S1-head-up state. The domain closure is mediated by double hydrogen bonds connecting ARG of ChainA with ASN and CYS residue of ChainB. B. Inter-chain RBD-S2 domain closure in the S1-head-down state. The S2 stalk connection with RBD is mediated by a proline residue of ChainA with the formation of a CH-п type interaction with tyrosine and hydrophobic interaction with another proline of ChainB.

**Fig. S2:**
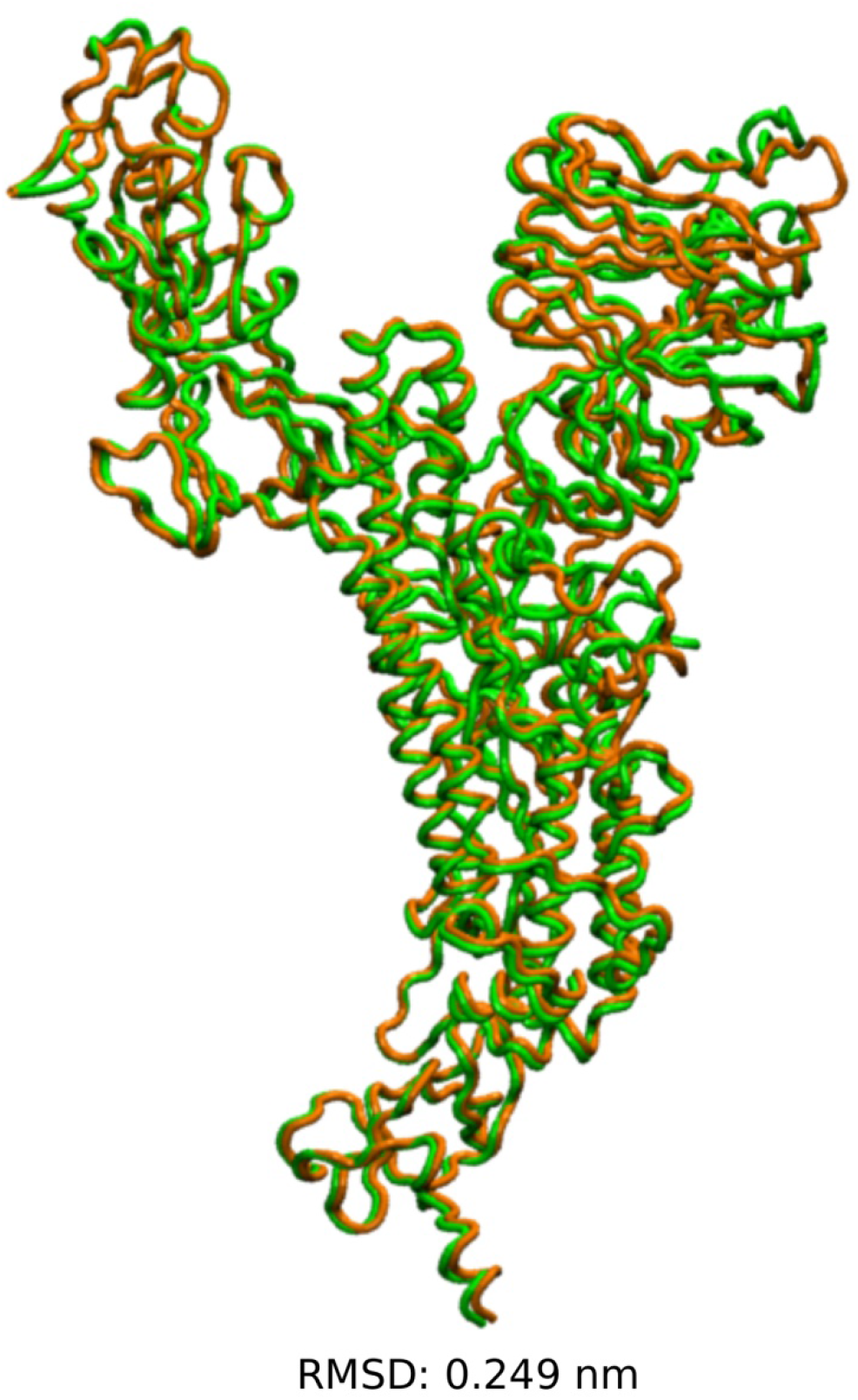
The structural alignment of two chains in the S1-head-down state. ChainB (orange) and ChainC (green) in the S1-head-down state extracted from the Cryo-EM structure (pdb:6vsb) of trimeric spike. Low RMSD between these two chains suggests that contact information extracted from any of these chains will be equivalent. This supports our contact map generation shown in the method pipeline.

**Fig. S3.**
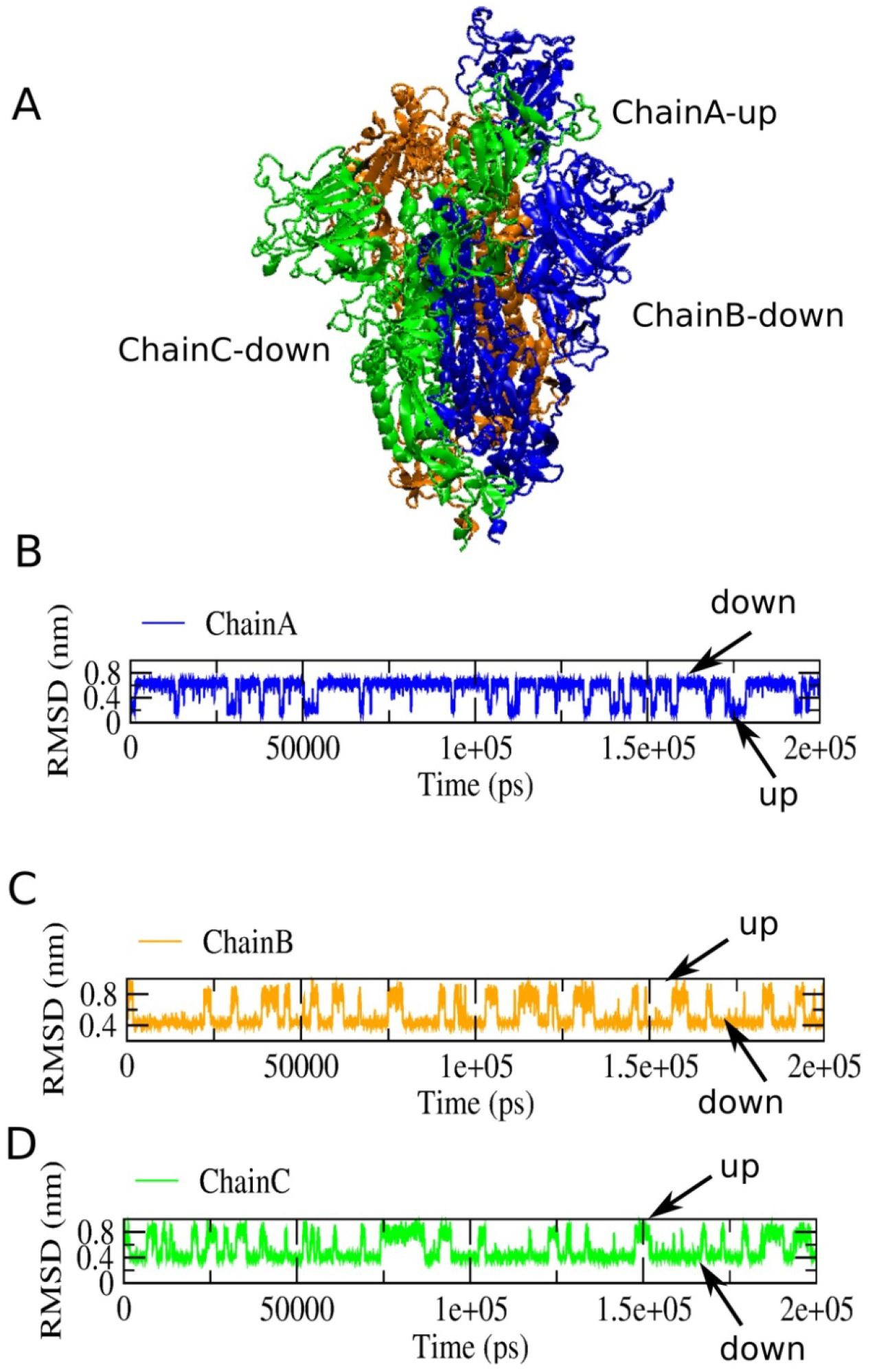
RMS deviation of each chain from their initial state during a typical simulation progress. A. The initial state of Chain A in the trimeric spike was in ‘S1-head-up’ state and Chain B/C was in ‘S1-head-down’ state. B. The lower RMSD for Chain A corresponds to Chain A’s head-up state. C. The lower RMSD for chain B corresponds to Chain B ‘s head-down state. D. The lower RMSD for Chain C corresponds to Chain C’s head-down state. The RMSD analyses ensure the correctness of the simulation progress and the emergence of the correct structure.

**Fig. S4.**
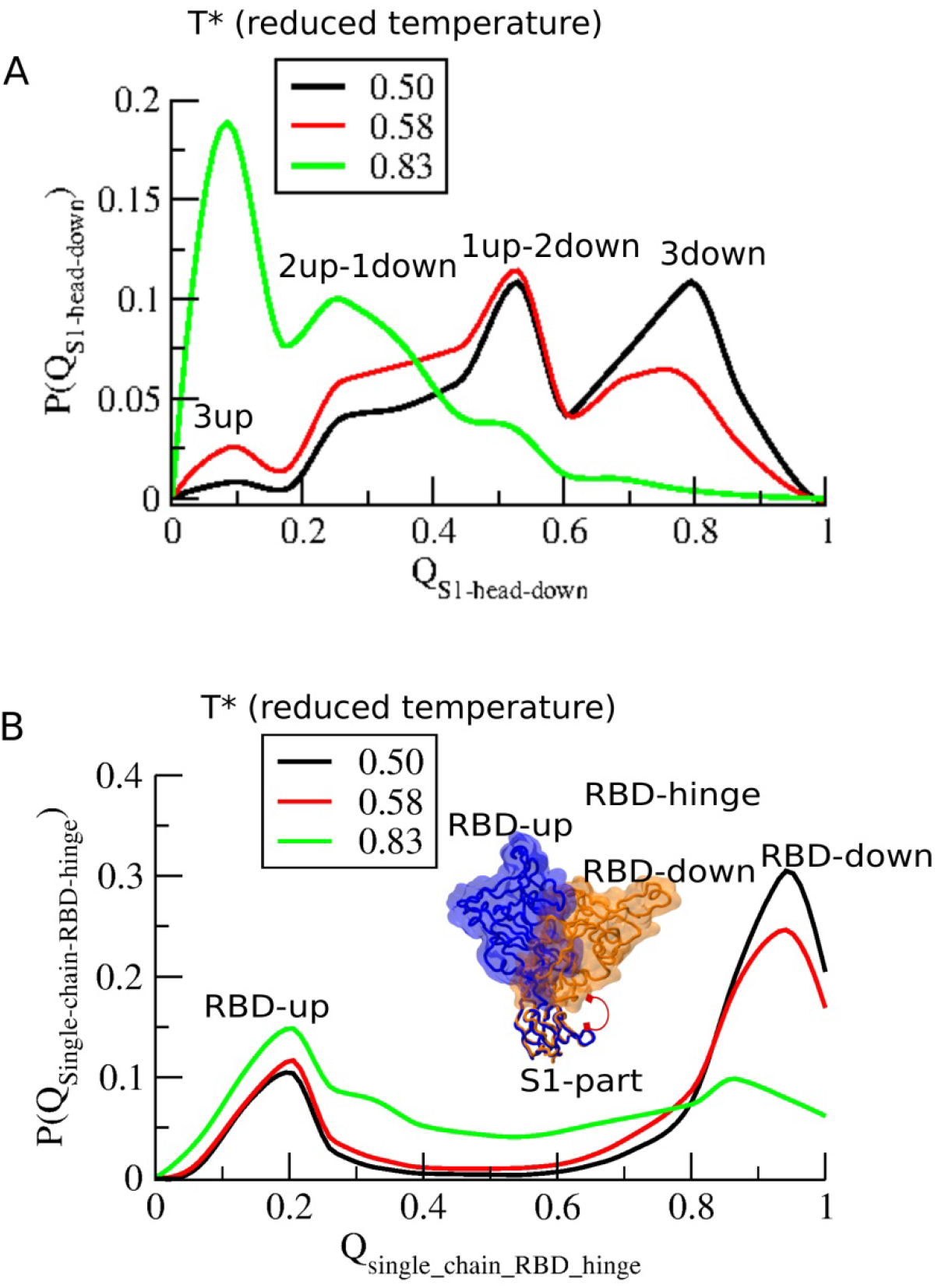
Temperature dependence of S1-head up-down transition and RBD open-close breathing transition. A. Population distribution as a function of the of native inter-chain contacts formed in the S1-head-down state as shown in Fig. S1. Four states emerge as shown in Fig. 3B. As temperature increases the population shifts more towards the S1-head-up state conformations indicating that S1-head-up states are more dynamic and entropically stable.*******Note that the dynamical transition between 1up-2down and 2up-1down states may tolerate a wide range of temperatures by a population shift mechanism.******* B. Population distribution as a function of the fraction of native intra-chain hinge-region contacts formed by the RBD. A bimodal distribution reflects the ‘RBD-up’ and the ‘RBD-down’ states for any individual chain being in trimeric spike. As the temperature increases, RBD-up started populating more. Temperature analysis helps to choose an intermediate temperature to obtain correct population distribution.

**Fig. S5.**
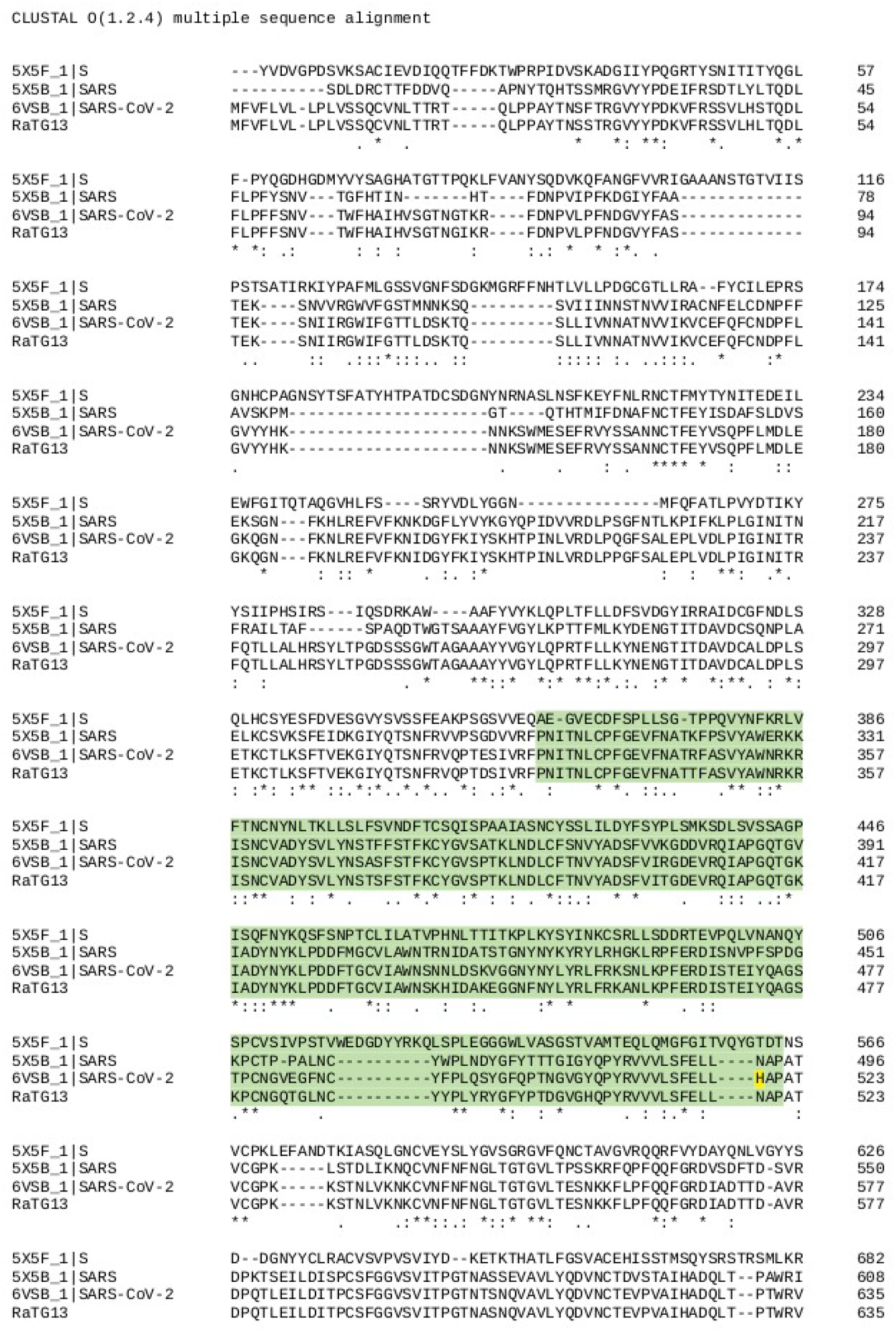
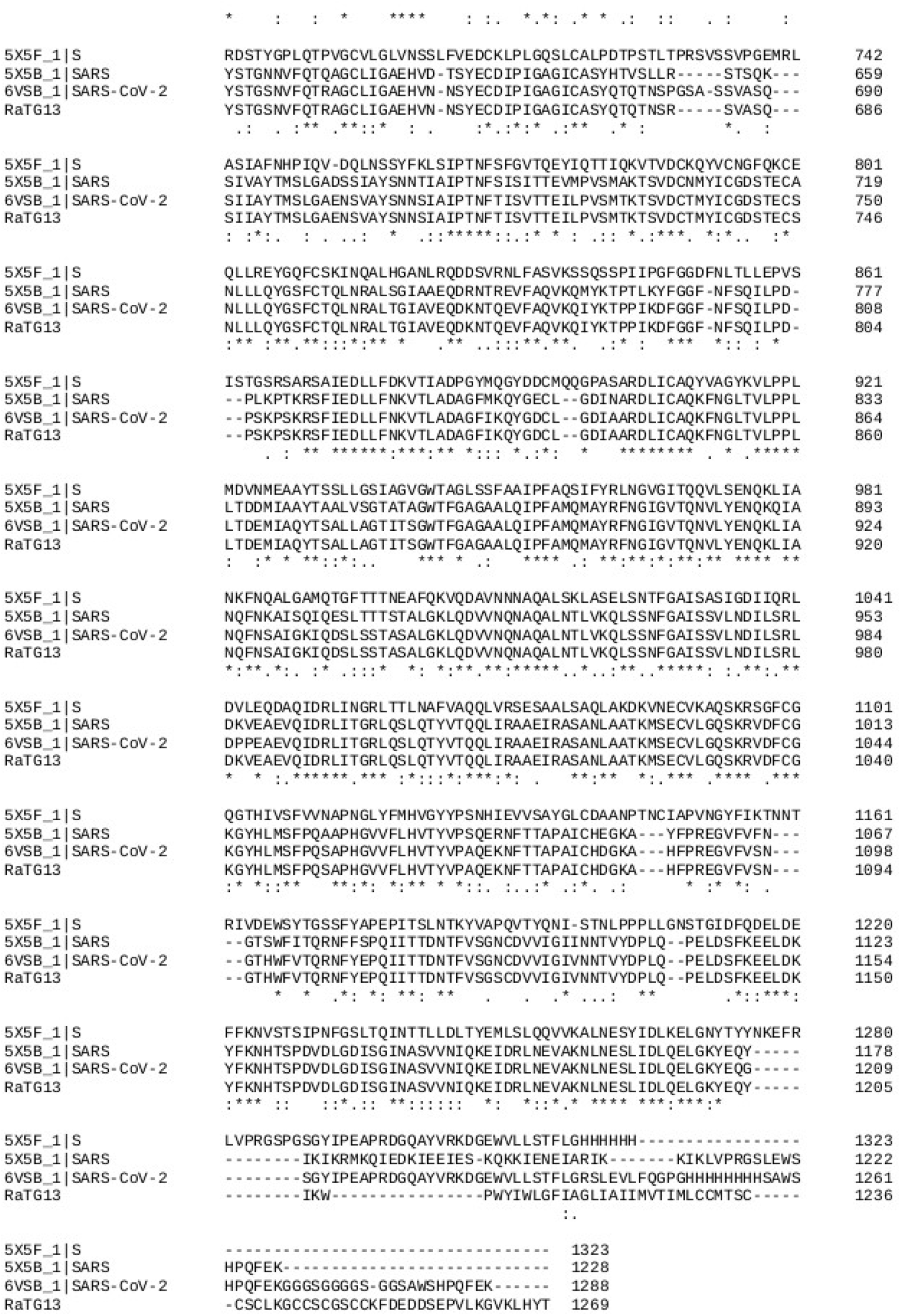
Sequence alignment of SARS-CoV-2 spike (pdb: 6vsb) with that of SARS-CoV spike(pdb:5×5b), MERS-CoV spike (pdb: 5×5f) and RaTG13 spike. Only the RBD is highlighted in green. The unique histidine residue (highlighted in yellow) of the RBD of SARS-CoV-2 is noted. Identical residues are denoted by an “*” beneath the consensus position. The multiple sequence alignment is continued over the next page.

**Fig. S6.**
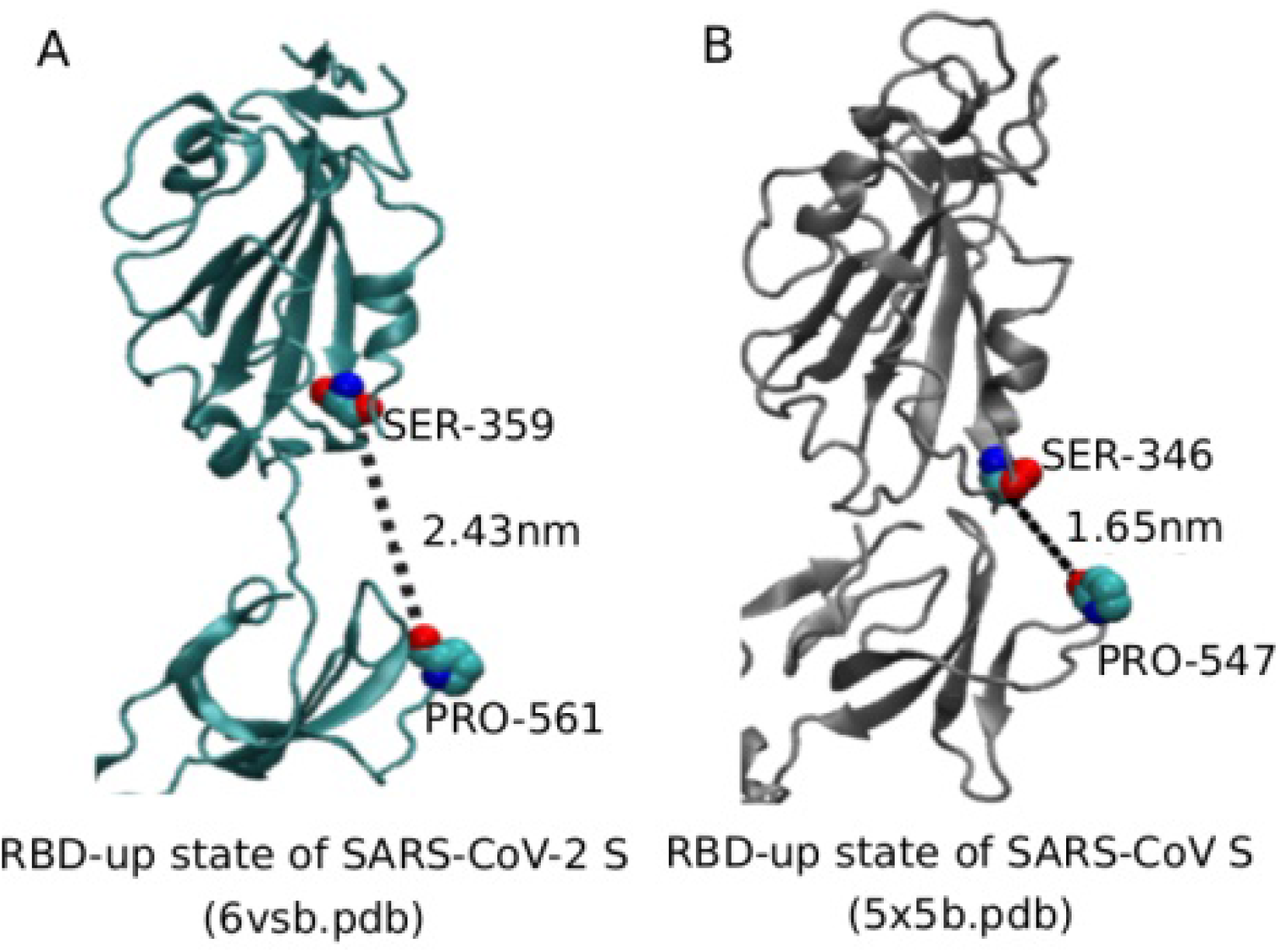
The opening of RBD-S1 cleft in the ‘S1-head-up’ state of SARS-CoV-2 S differs from that of SARS-CoV S. The opening is measured by a characteristic distance between a serine and proline residues at two edges of the cleft. For SARS-CoV-2 S the distance is 2.43 nm while for SARS-CoV-S, it is 1.65nm. It appears that inter-chain RBD-NTD connection influences the SARS-CoV S RBD hinge motion significantly where the cleft opening is supported by those inter-chain interactions.

**Fig. S7.**
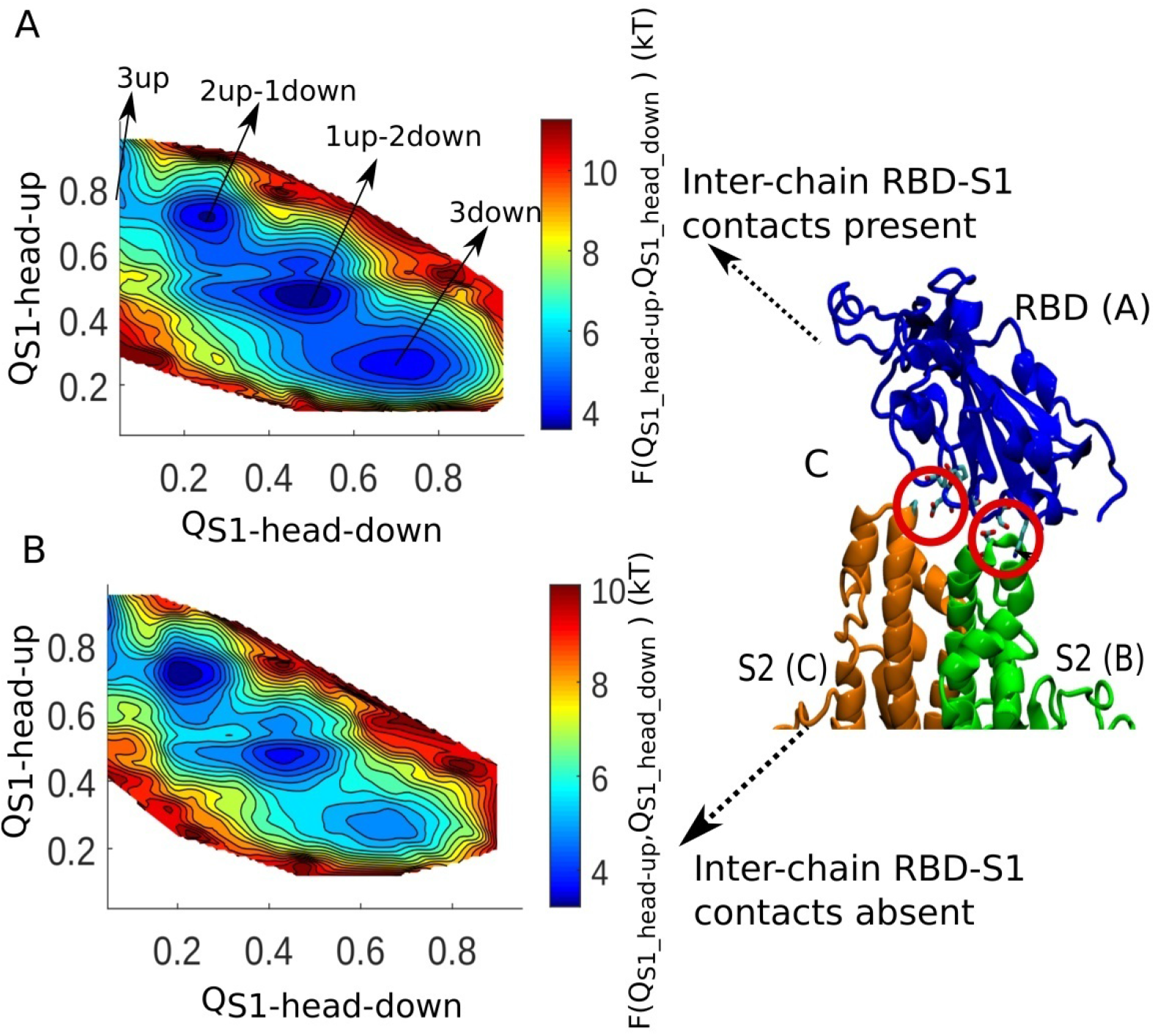
The free energy landscape in the presence and absence of inter-chain RBD-S1 contacts. A. In the presence of inter-chain RBD-S1 contacts, the enhanced population of the 1up-2down compared to 2up-1down. B. In the absence of inter-chain RBD-S1 contacts, the population shifts from 1up-2down state 2up-1down state.

**Movie S1: Conformational dynamics of full-length trimeric SARS-CoV-2 spike protein showing rapid symmetry breaking.**

**Movie S2:Conformational dynamics of full-length trimeric SARS-CoV-2 spike protein showing rapid symmetry breaking. In this movie the NTD domains are not shown for better demonstration of the RBD movement.**

**Movie S3: Conformational dynamics of a monomer of the full-length SARS-CoV-2 showing RBD hinge motion.**

## References and Notes

1. in Microbial Evolution and Co-Adaptation: A Tribute to the Life and Scientific Legacies of Joshua Lederberg: Workshop Summary. (Washington (DC), 2009).

2. W. H. McNeill, Plagues and peoples. (Anchor Books, New York, 1998), pp. 365 p.

3. J. Zheng, SARS-CoV-2: an Emerging Coronavirus that Causes a Global Threat. International journal of biological sciences 16, 1678 (2020).

4. X. Xu et al., Evolution of the novel coronavirus from the ongoing Wuhan outbreak and modeling of its spike protein for risk of human transmission. Science China. Life sciences 63, 457 (Mar, 2020).

5. T. T. Lam et al., Identifying SARS-CoV-2 related coronaviruses in Malayan pangolins. Nature, (Mar 26, 2020).

6. F. Li, Structure, Function, and Evolution of Coronavirus Spike Proteins. Annual review of virology 3, 237 (Sep 29, 2016).

7. D. Wrapp et al., Cryo-EM structure of the 2019-nCoV spike in the prefusion conformation. Science 367, 1260 (Mar 13, 2020).

8. A. C. Walls et al., Structure, Function, and Antigenicity of the SARS-CoV-2 Spike Glycoprotein. Cell, (Mar 6, 2020).

9. R. Yan et al., Structural basis for the recognition of SARS-CoV-2 by full-length human ACE2. Science 367, 1444 (Mar 27, 2020).

10. M. Gui et al., Cryo-electron microscopy structures of the SARS-CoV spike glycoprotein reveal a prerequisite conformational state for receptor binding. Cell research 27, 119 (Jan, 2017).

11. D. Wrapp, J. S. McLellan, The 3.1-Angstrom Cryo-electron Microscopy Structure of the Porcine Epidemic Diarrhea Virus Spike Protein in the Prefusion Conformation. Journal of virology 93, (Dec 1, 2019).

12. Y. Yuan et al., Cryo-EM structures of MERS-CoV and SARS-CoV spike glycoproteins reveal the dynamic receptor binding domains. Nature communications 8, 15092 (Apr 10, 2017).

13. M. Hoffmann et al., SARS-CoV-2 Cell Entry Depends on ACE2 and TMPRSS2 and Is Blocked by a Clinically Proven Protease Inhibitor. Cell, (Mar 4, 2020).

14. M. Yuan et al., A highly conserved cryptic epitope in the receptor-binding domains of SARS-CoV-2 and SARS-CoV. Science, (Apr 3, 2020).

15. C. Clementi, H. Nymeyer, J. N. Onuchic, Topological and energetic factors: what determines the structural details of the transition state ensemble and “en-route” intermediates for protein folding? An investigation for small globular proteins. Journal of molecular biology 298, 937 (May 19, 2000).

16. A. Waterhouse et al., SWISS-MODEL: homology modelling of protein structures and complexes. Nucleic acids research 46, W296 (Jul 2, 2018).

17. Z. Li et al., The human coronavirus HCoV-229E S-protein structure and receptor binding. eLife 8, (Oct 25, 2019).

18. J. K. Noel et al., SMOG 2: A Versatile Software Package for Generating Structure-Based Models. PLoS computational biology 12, e1004794 (Mar, 2016).

19. J. K. Noel, P. C. Whitford, J. N. Onuchic, The shadow map: a general contact definition for capturing the dynamics of biomolecular folding and function. The journal of physical chemistry. B 116, 8692 (Jul 26, 2012).

20. P. E. Leopold, M. Montal, J. N. Onuchic, Protein folding funnels: a kinetic approach to the sequence-structure relationship. Proceedings of the National Academy of Sciences of the United States of America 89, 8721 (Sep 15, 1992).

21. P. G. Wolynes, Symmetry and the energy landscapes of biomolecules. Proceedings of the National Academy of Sciences of the United States of America 93, 14249 (Dec 10, 1996).

22. P. G. Wolynes, W. A. Eaton, A. R. Fersht, Chemical physics of protein folding. Proceedings of the National Academy of Sciences of the United States of America 109, 17770 (Oct 30, 2012).

23. R. Zwanzig, A. Szabo, B. Bagchi, Levinthal’s paradox. Proceedings of the National Academy of Sciences of the United States of America 89, 20 (Jan 1, 1992).

24. J. D. Bryngelson, J. N. Onuchic, N. D. Socci, P. G. Wolynes, Funnels, pathways, and the energy landscape of protein folding: a synthesis. Proteins 21, 167 (Mar, 1995).

25. B. Jana, C. Hyeon, J. N. Onuchic, The origin of minus-end directionality and mechanochemistry of Ncd motors. PLoS computational biology 8, e1002783 (2012).

26. X. Lin et al., Order and disorder control the functional rearrangement of influenza hemagglutinin. Proceedings of the National Academy of Sciences of the United States of America 111, 12049 (Aug 19, 2014).

27. H. Nymeyer, N. D. Socci, J. N. Onuchic, Landscape approaches for determining the ensemble of folding transition states: success and failure hinge on the degree of frustration. Proceedings of the National Academy of Sciences of the United States of America 97, 634 (Jan 18, 2000).

28. M. Brandl, M. S. Weiss, A. Jabs, J. Suhnel, R. Hilgenfeld, C-H...pi-interactions in proteins. Journal of molecular biology 307, 357 (Mar 16, 2001).

29. F. Madeira et al., The EMBL-EBI search and sequence analysis tools APIs in 2019. Nucleic acids research 47, W636 (Jul 2, 2019).

30. R. N. Kirchdoerfer et al., Stabilized coronavirus spikes are resistant to conformational changes induced by receptor recognition or proteolysis. Scientific reports 8, 15701 (Oct 24, 2018).

## References and Notes

2. W. H. McNeill, Plagues and peoples. (Anchor Books, New York, 1998), pp. 365 p.

31. M. Dutta, B. Jana, Role of AAA3 Domain in Allosteric Communication of Dynein Motor Proteins. ACS omega 4, 21921 (Dec 24, 2019).

32. C. Ghosh, B. Jana, Intersubunit Assisted Folding of DNA Binding Domains in Dimeric Catabolite Activator Protein. The journal of physical chemistry. B 124, 1411 (Feb 27, 2020).

33. N. V. Dokholyan, SpringerLink (Online service), Computational Modeling of Biological Systems : From Molecules to Pathways. Biological and Medical Physics, Biomedical Engineering, , pp. 1 online resource (VI, 366 p.).

34. C. L. Brooks, 3rd, M. Gruebele, J. N. Onuchic, P. G. Wolynes, Chemical physics of protein folding. Proceedings of the National Academy of Sciences of the United States of America 95, 11037 (Sep 15, 1998).

35. K. A. Dill, H. S. Chan, From Levinthal to pathways to funnels. Nature structural biology 4, 10 (Jan, 1997).

36. B. Jana, F. Morcos, J. N. Onuchic, From structure to function: the convergence of structure based models and co-evolutionary information. Physical chemistry chemical physics : PCCP 16, 6496 (Apr 14, 2014).

37. N. D. Socci, J. N. Onuchic, P. G. Wolynes, Protein folding mechanisms and the multidimensional folding funnel. Proteins 32, 136 (Aug 1, 1998).

38. P. G. Wolynes, J. N. Onuchic, D. Thirumalai, Navigating the folding routes. Science 267, 1619 (Mar 17, 1995).

39. N. R. Eddy, J. N. Onuchic, Rotation-Activated and Cooperative Zipping Characterize Class I Viral Fusion Protein Dynamics. Biophysical journal 114, 1878 (Apr 24, 2018).

40. J. D. Honeycutt, D. Thirumalai, The nature of folded states of globular proteins. Biopolymers 32, 695 (Jun, 1992).

41. B. Jana, J. N. Onuchic, Strain Mediated Adaptation Is Key for Myosin Mechanochemistry: Discovering General Rules for Motor Activity. PLoS computational biology 12, e1005035 (Aug, 2016).

